# Preservation of intracellular protein-protein interactions in archived cytological specimens: Optimization of preanalytical variables for in situ proximity ligation assay

**DOI:** 10.64898/2026.10.06.757074

**Authors:** Caroline Maria Dahlström, Jan-Michaél Hirsch, Liza Löf

**Affiliations:** Molecular tools and functional genomics, Department of Immunology, Genetics and Pathology, Uppsala University, Sweden; Department of Surgical Sciences, Oral & Maxillofacial Surgery, Uppsala University, Sweden; Folktandvården Stockholms län AB, Stockholm, Sweden

**Keywords:** cytology, isPLA, storage, protein-protein interactions, molecular biology

## Abstract

**Context:** Cytological specimens allow minimally invasive, repeated sampling for diagnosis and molecular testing. However, cytology biobanking protocols have primarily been optimized for morphology, and the effects of preanalytical handling and storage on intracellular proteins remain unclear.

**Objective:** To assess how fixation, staining, storage temperature, and duration affect intracellular protein preservation in cytological specimens and to identify conditions suitable for in situ proximity ligation assay (isPLA).

**Design:** Human cancer cell lines were processed under different preanalytical conditions. Intracellular proteins and protein–protein interactions were assessed by isPLA, fluorescence microscopy, and quantitative image analysis. Slides were stored at room temperature or −20°C, with or without May–Grünwald–Giemsa (MGG) staining.

**Results:** Storage temperature had the greatest effect on protein signal. Unstained slides stored at −20°C retained strong signals for up to 24 months, whereas room temperature storage reduced detectable protein interactions by approximately 95%. MGG staining further reduced signal intensity during prolonged storage. Nevertheless, stained slides remained suitable for isPLA when staining, microscopic evaluation, destaining, and antibody labeling were completed on the same day. Prolonged room temperature storage of stained slides caused loss of antigenicity, increased autofluorescence, and false negative findings.

**Conclusions:** Low temperature storage and standardized preanalytical handling preserve intracellular protein signals in cytological specimens. These conditions support reliable single cell protein profiling of archived samples and their use in biomarker research and translational pathology.

## 1.1 Introduction

Biobanking plays a crucial role in biomedical research and the advancement of personalized medicine. Reliable preservation of biological specimens is essential for accurate biomarker analyses and retrospective molecular studies ^1–3^. Cytological samples are typically processed immediately after collection and they are seldom stored for extended periods prior to analysis^4,5^. When archived in biobanks nucleic acids such as DNA and RNA are generally well preserved under routine cytological storage conditions and therefore widely used for genomic and transcriptomic investigations ^6–8^. In contrast, intracellular proteins are particular susceptible to pre-analytical factors, including fixation, processing, and storage conditions, which may compromise protein integrity through degradation, conformational alterations, and epitope masking ^9–11^. Because proteins represent the functional products of gene expression and cellular signaling, preservation of protein integrity is essential for reliable biomarker discovery, molecular characterization, and translational research.

Cytology is a rapid, minimally invasive diagnostic technique that enables the evaluation of evaluating large numbers of cells using light microscopy. Liquid-based cytology (LBC) methods are well established in gynaecological cytology and have increasingly been applied to oral cytology and other non-gynecological specimens because they provide standardized specimen preparation, improved cellular preservation, and cleaner microscopic backgrounds^5^. Although these preparations provide excellent cellular morphology, they have primarily been optimized for cytomorphological assessment rather than for molecular analyses.

Consequently, the potential of cytological specimens for intracellular protein profiling and other single cell molecular applications remains insufficiently explored.

In routine diagnostic practice, previously stained cells may be destained and subsequently processed for immunocytochemistry (ICC), enabling additional immunophenotypic characterization while preserving valuable cellular material ^12–14^.

However, the suitability of long-term archived cytology specimens for protein-based analyses has received limited attention. In particular, the effects of prolonged storage on protein integrity, antigen preservation, and epitope accessibility remain poorly characterized, limiting the potential use of archived cytology material for retrospective biomarker studies.

While conventional ICC serves to identify the distribution of proteins in a sample, in situ proximity ligation assay (isPLA) serves to enhance sensitive detection of proteins, as each target must be recognized by a pair of antibodies targeting specific epitopes^15^. The requirement for dual recognition also permits analyses of protein-protein interactions and post-translational modifications within cells. The assays employ pairs of oligonucleotide-conjugated antibodies that guide the formation of circular DNA strands upon their proximal binding. The DNA circles are allowed to template locally amplified DNA products via rolling-circle amplification (RCA). Fluorophore-labelled probes are then hybridized to the amplified RCA products, enabling highly specific and sensitive visualization of the protein targets of interest ^15–17^. isPLA depends on preservation of protein conformation and epitope accessibility by the antibody reagents and, pre-analytical handling is therefore expected to have a major influence on assay performance.

To investigate these factors, established human cancer cell lines were used as standardized experimental models to systematically evaluate the effects of cytological preparation, fixation, staining, storage temperature, and storage duration on intracellular protein preservation. The use of well-characterized cell lines enabled reproducible assessment of pre-analytical variables under controlled experimental conditions.

### 1.2 Aim

This study aimed to determine how pre-analytical factors, including fixation, staining procedures, storage temperature, and storage duration, influence intracellular protein integrity and the detection of protein-protein interactions in cytological specimens. Furthermore, we sought to establish optimized handling and storage protocols that preserve protein integrity and enable reliable downstream molecular analyses using *is*PLA, thereby expanding the utility of archived cytological specimens for biomarker discovery, translational research, and precision medicine.

## 2. Materials and Methods

### 2.1 Cell Lines

Human HaCaT keratinocytes, Jurkat T-cell leukemia cells, GM12878 lymphoblastoid B cells, and SKBR-3 breast cancer cells were obtained from the American Type Culture Collection (ATCC). Jurkat, GM12878 and SKBR-3 cells were all cultured in RPMI 1640 medium, where HaCaT cells maintained in Dulbecco’s Modified Eagle Medium (DMEM). All culture media were supplemented with 10% fetal bovine serum (FBS), 2 mM L-glutamine, and 100 U/mL–100 µg/mL penicillin-streptomycin (Sigma-Aldrich, USA). The cultures were incubated at 37°C in a humidified atmosphere with 5% CO□.

### 2.2 Cell Preparation and Fixation

The non-adherent GM12878 and Jurkat cells were harvested, centrifuged at 130 × g for 2 minutes, and the medium supernatant was discarded. The adherent HaCaT cells were detached using 0,25% Trypsin-EDTA (Cat. No. T4049, Sigma-Aldrich, USA) and followed by pelleting via centrifugation at 130 x g for 2 minutes; the supernatant of the medium was discarded. The cells were fixed in 99% methanol, and squash cell smears were prepared by placing a 20 µL drop of the cell suspension between two glass slides and gently compressing to distribute the cells into a monolayer. Slides were air-dried and subsequently processed according to the experimental protocol, including staining and storage at either room temperature (RT) or - 20°C.

Adherent SKBR-3 cells were first treated and detached using 0.25% (w/v) trypsin-EDTA and resuspended in complete culture medium, and seeded onto Millicell EZ SLIDE 8-well glass(Millipore, USA), After incubation for 24 hours at 37°C, the cells were washed twice with 1× PBS, fixed in 1% formaldehyde (PFA) diluted in PBS, and subsequently stored at - 20°C for periods ranging from 1 to 12 months before analysis.

### 2.3 Staining procedure with May-Grunwald-Giemsa (MGG)-staining

GM12878 cytology cell slides were stained using the MGG-protocol. Slides were immersed in May–Grünwald solution (Sigma-Aldrich, USA) for 5 minutes, rinsed in PBS, pH 7.4 and subsequently stained in 5% Giemsa solution (Sigma-Aldrich, USA) for 5 minutes. The slides were washed in tap water and air-dried before storage in RT. Unless otherwise specified, no destaining was performed before antibody incubation. Study design in Figure 1 and the layout of conditions in Table 1.

**Figure 1.**
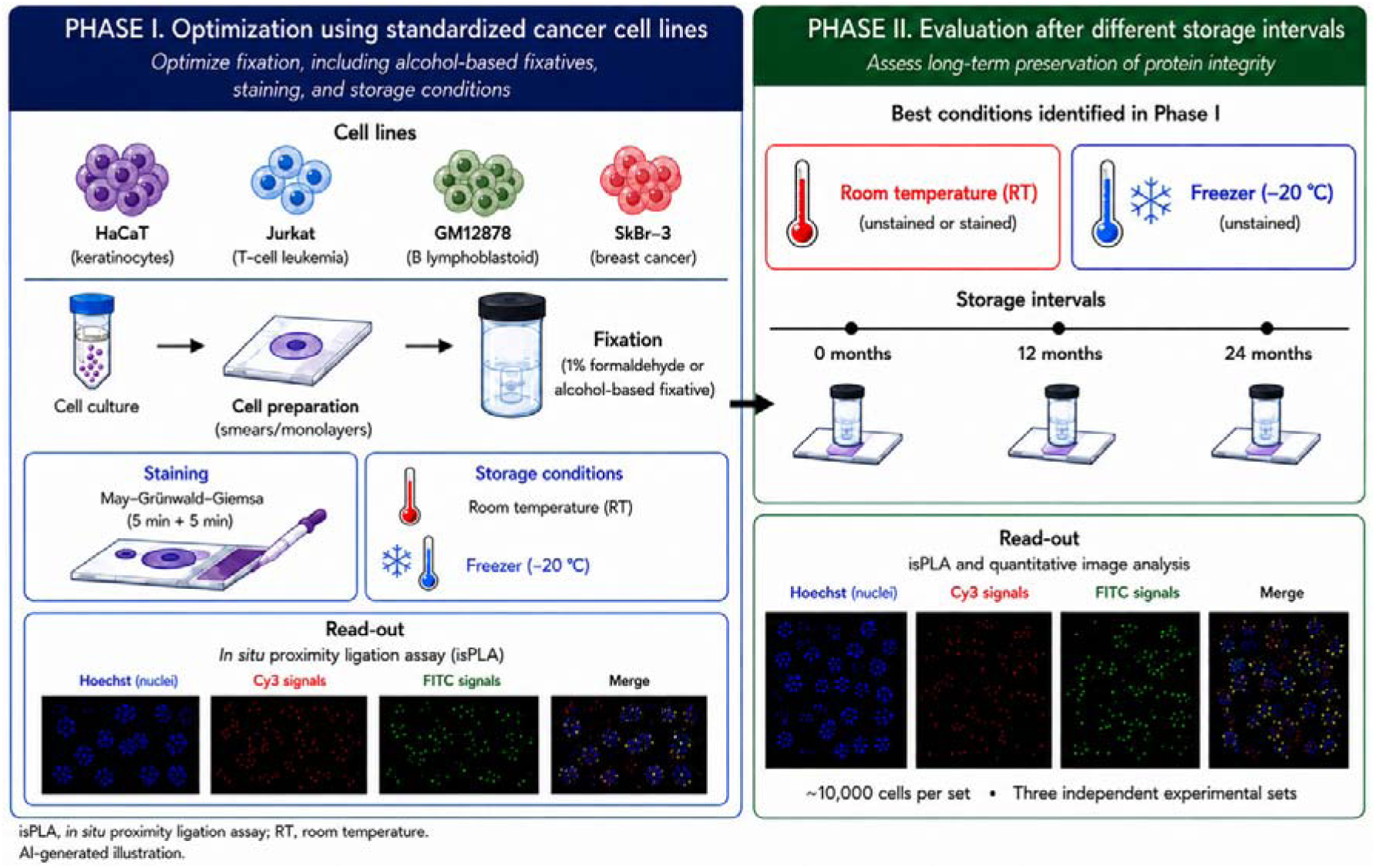
Overview of the experimental study design. The study was conducted in two sequential phases. In phase I, four standardized cancer cell lines(HaCaT, Jurkat, GM12878, and SKBR-3) were used to evaluate preanalytical conditions, including formaldehyde- or alcohol-based fixation, MGG, and storage conditions (RT or -20°C). In phase II, stained and unstained cytological preparations were stored at room temperature or -20°C for different intervals to evaluate the effects of storage temperature and duration on intracellular protein preservation. Protein expression and protein-protein interactions were assessed by isPLA, followed by fluorescence microscopy and quantitative image analysis. Approximately 10,000 cells were analysed per condition in three independent experimental sets.

**Table 1.** Cell lines and preanalytical conditions evaluated in the study. . SKBR-3 cells were fixed with 1% formaldehyde, whereas GM12878, Jurkat, and HaCaT cells were fixed with 99% methanol. Slides were analyzed after storage at RT or -20°C, with or without prior MGG staining. Baseline indicates newly prepared slides analyzed without prolonged storage. MGG: May-Grunwald-Giemsa, RT: room temperature

| *Cell line type* | *Cell line* | *MGG stain* | *Fixation* | *Storage temp* | *Time before analysis* |
| --- | --- | --- | --- | --- | --- |
| *Hematologic* | GM12878 | No | 99% methanol | -20 °C | 24 months |
| *Hematologic* | GM12878 | Yes | 99% methanol | RT | 24 months |
| *Hematologic* | Jurkat | No | 99% methanol | RT | 24 months |
| *Epithelial* | HaCaT | No | 99% methanol | RT | 24 months |
| *Epithelial* | SKBR-3 | No | 1% PFA | -20 °C | 0 months |
| *Epithelial* | SKBR-3 | No | 1% PFA | -20 °C | 12 months |

### 2.4 Permeabilization, Blocking, and isPLA

Slides were equilibrated to RT, permeabilized by adding 40 µL of 0.25% Triton X-100 (Sigma-Aldrich, St. Louis, USA) for 5 min at RT, and blocked with Odyssey Blocking Buffer ( LI-COR, USA) in humid chamber at 37°C for 30 min.

Primary antibodies, MEK (MAB2079, Bio-Techne), ERK (AF1230, Bio-Techne), Akt3 (MAB14631, Bio-Techne), and pan-Akt (MAB2055, Bio-Techne), conjugated to oligonucleotides and hybridized to padlock-probes, were diluted to a final concentration of 5 µg/mL in blocking buffer and added to the slides and incubated in a humid chamber at 37°C for 1 hour. Following washing, ligation and RCA were performed according to standard isPLA protocol. The ligation mix, consisting of 0.25 mg/mL bovine serum albumin (BSA) (New England Biolabs, UK), 0.5% Tween-20, 0.5 mg/mL NaCl, 0.5 mM ATP(Thermo Fisher Scientific, USA), 0.05 U/µL T4 DNA ligase(Thermo Fisher Scientific), and 1 U/µL UNG enzyme(Thermo Fisher Scientific) diluted in nuclease-free ddH O, was added to each well and incubated at 37°C for 30 min in a humidified chamber. The RCA mix, containing 0.25 mg/mL BSA( New England Biolabs,UK), 2× Phi29 DNA polymerase buffer, 0.25 mM dNTPs, 0.25 U/µL Phi29 DNA polymerase from Termo Fisher Scientific, USA. 0.01 µM detection oligonucleotides carrying either ATTO 550 or ATTO 488flourophores from IdT, USA, and 0.25 µM Hoechst (Thermo Fisher Scientific, USA), all diluted in nuclease-free ddH O, was added to each well. Slides were incubated at 37°C for 60 min in a humid chamber. After RCA, the slides were washed in 1xTBS-0.05% Tween 20 under gentle shaking for 10 min.

Finally, the slides were mounted using ProLong™ Gold Antifade Mountant (Thermo Fisher Scientific, USA).

### 2.5 Microscopy

Slides were imaged using fluorescence microscopy with a Zeiss Imager Z2 microscope equipped with a Hamamatsu C11440 camera and controlled by Zen 2 Blue Edition software. Samples were excited using an HXP 120 V light source, and images were acquired using a 20× apochromatic objective lens, 0.8 numerical aperture. No immersion medium was used. Instrument calibration and manual focusing were performed before image acquisition.

### 2.6 Image Analysis

Image analysis was performed using CellProfiler (Broad Institute,MA, USA). Image analysis was performed using a customized CellProfiler pipeline. Nuclei were identified based on Hoechst staining and used as the basis for subsequent segmentation of individual cells. Background signals were reduced prior to object detection, and objects were filtered according to predefined size and fluorescence intensity thresholds to minimize the inclusion of non-specific signals and imaging artifacts. isPLA signals were subsequently detected and quantified as discrete fluorescent dots in the Cy3 and FITC channels.

A total of 39,177 cells derived from five independently prepared experimental slide sets were analyzed. Quantitative comparisons were based on the means of biological replicates rather than individual cells for each experimental set under three independent experimental series comprising stained and unstained slides stored at RT or -20°C. The number of isPLA signals per cell and fluorescence channel was quantified and averaged across replicates. A separate CellProfiler pipeline was optimized for the SKBR-3 dataset to account for differences in noise level, background noise and autofluorescence, cell morphology, including acquisition settings, and the objects mentioned above.

### 2.7 Experimental design and statistical analysis

All experiments were performed in three independent biological replicates. Within each biological replicate, cytological preparations were generated under identical experimental conditions and analyzed using standardized image acquisition settings. Approximately 10,000 cells were quantified per experimental condition by automated CellProfiler analysis. Cell-level measurements were aggregated within each biological replicate before statistical analysis.

Biological replicates consisted of independently cultured and processed cell preparations performed on separate occasions. Technical replicates consisted of repeated image acquisitions and CellProfiler analyses performed using identical acquisition settings.

This controlled experimental study was designed primarily to evaluate the magnitude and consistency of pre-analytical effects rather than to test formal statistical hypotheses.

Quantitative measurements from independent experiments demonstrated highly consistent trends across all experimental conditions.

### 2.8 Ethical requirements

The study was conducted in accordance with the Declaration of Helsinki. The use of established human cell lines did not require ethical approval.

## 3. Results

The primary objective of this study was to determine whether intracellular proteins and protein-protein interactions could be preserved in cytological specimens during storage. Protein interactions were visualized using isPLA, imaged by widefield fluorescence microscopy, and quantified using Cellprofiler software.

### 3.1 Storage temperature markedly influences intracellular protein preservation

A total of 39,177 cells were analyzed from five independent experimental slide sets. GM12878 cells stored at -20°C served as the reference condition and were considered to represent complete preservation of detectable protein signals (100%). The results of the loss of signals, are shown in relative numbers to the optimal conditions. The initial results are signals per cell, and the signals per cell are converted into to loss of percentage. The optimal conditions assumed to show 100% potential signals, which are non-MGG stained cells stored in -20**°**C. GM12878 slides stored at RT, either unstained or after MGG staining, demonstrated a marked reduction in detectable isPLA signals. The overall signal frequency decreased by 94-99%(mean difference 95.2;95% Cl 92.4-97.8%;P < 0.001), indicating substantial deterioration of intracellular protein integrity during RT-storage (Figure 2).

**Figure 2.**
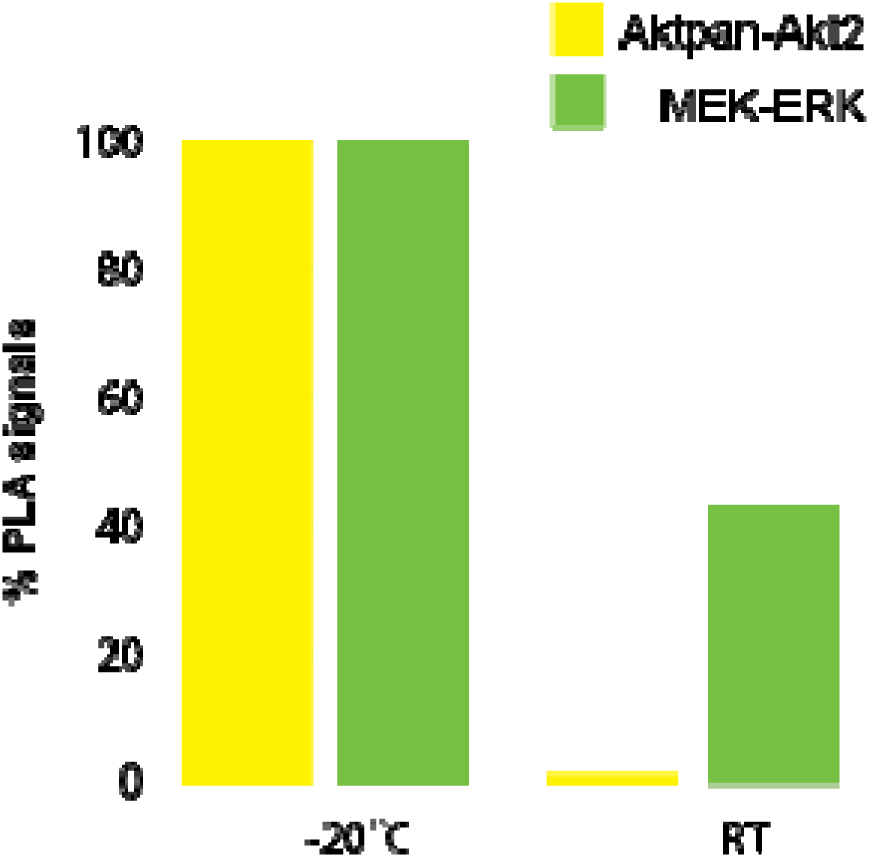
Effect of storage temperature on preservation of intracellular protein-protein interactions. Relative isPLA signal levels for Aktpan-Akt2 and MEK-ERK interactions are shown in GM12878 cells stored at -20**°**C or RT and in HaCaT cells stored at RT. Signal levels were normalized to those detected in unstained GM12878 cells stored at -20**°**C, which were defined as the reference condition (100%). Slides stored at RT showed a marked reduction in detectable isPLA signals compared with the reference condition. RT indicates room temperature; isPLA, in situ proximity ligation assay.

### 3.2 Differential preservation of Aktpan-Akt2 and MEK-ERK protein interactions in hematologic cells and epithelial cells

In GM12878 hematologic cells, Aktpan-Akt2 and MEK-ERK interactions differed according to storage conditions. For cells stored at -20°C, Aktpan-Akt2 interaction generated 2.2-fold (95% Cl, 1.9-2.6) more detectable isPLA signals than the MEK-ERK interaction. Mean difference 61% (95% Cl, 49-72%). Further, it was evident that proteins were better preserved in unstained cells stored at -20°C compared to MGG-stained cells, (Figure. 3). A higher autofluorescence intensity, no quantification was performed only noted, was also identified in MGG-stained material compared with the non-MGG-stained material Figure 5. After one year of storage of SKBR-3 cells at -20°C, the number of detectable MEK-ERK signals decreased to approximately 33% of expected values, whereas Aktpan-Akt2 signals decreased to 56% of expected values, Figure. 4.

**Figure 3.**
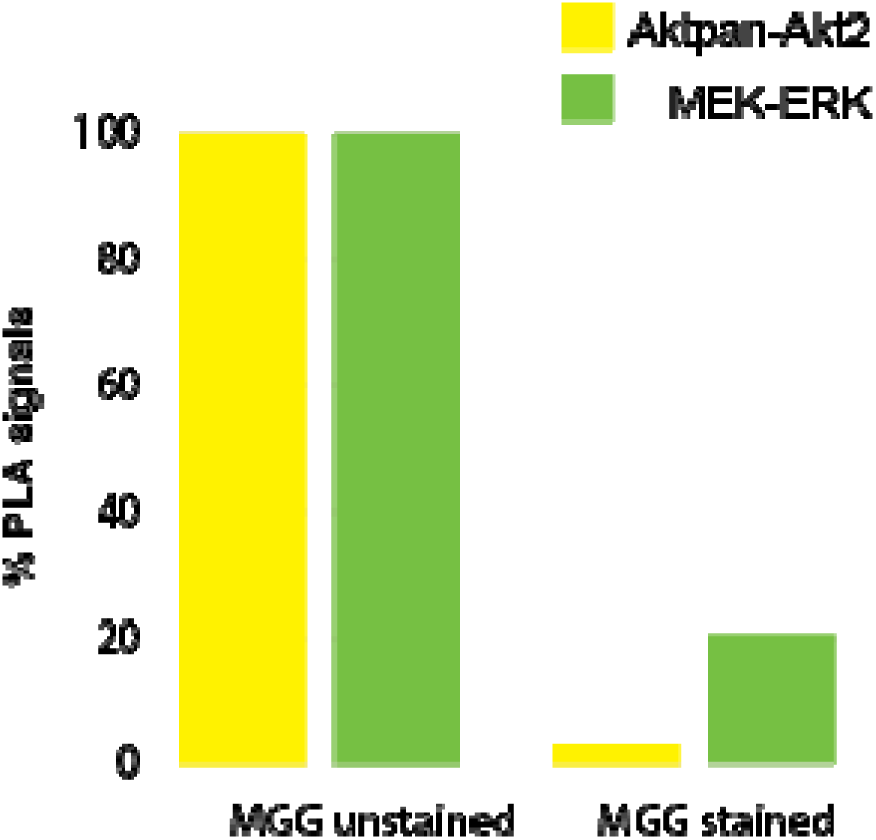
Effect of MGG staining and RT storage on detectable protein-protein interactions. Relative isPLA signal levels for the Aktpan-Akt2 and MEK-ERK interactions are shown for unstained GM12878 cells stored at -20**°**C and MGG-stained GM12878 cells stored at RT. Signal levels were normalized to those detected in unstained cells stored in - 20**°**C, which were defined as the reference condition (100%). MGG staining followed by RT storage resulted in a pronounced reduction in detectable isPLA signals. RT indicates room temperature; isPLA, in situ proximity ligation assay.

**Figure 4.**
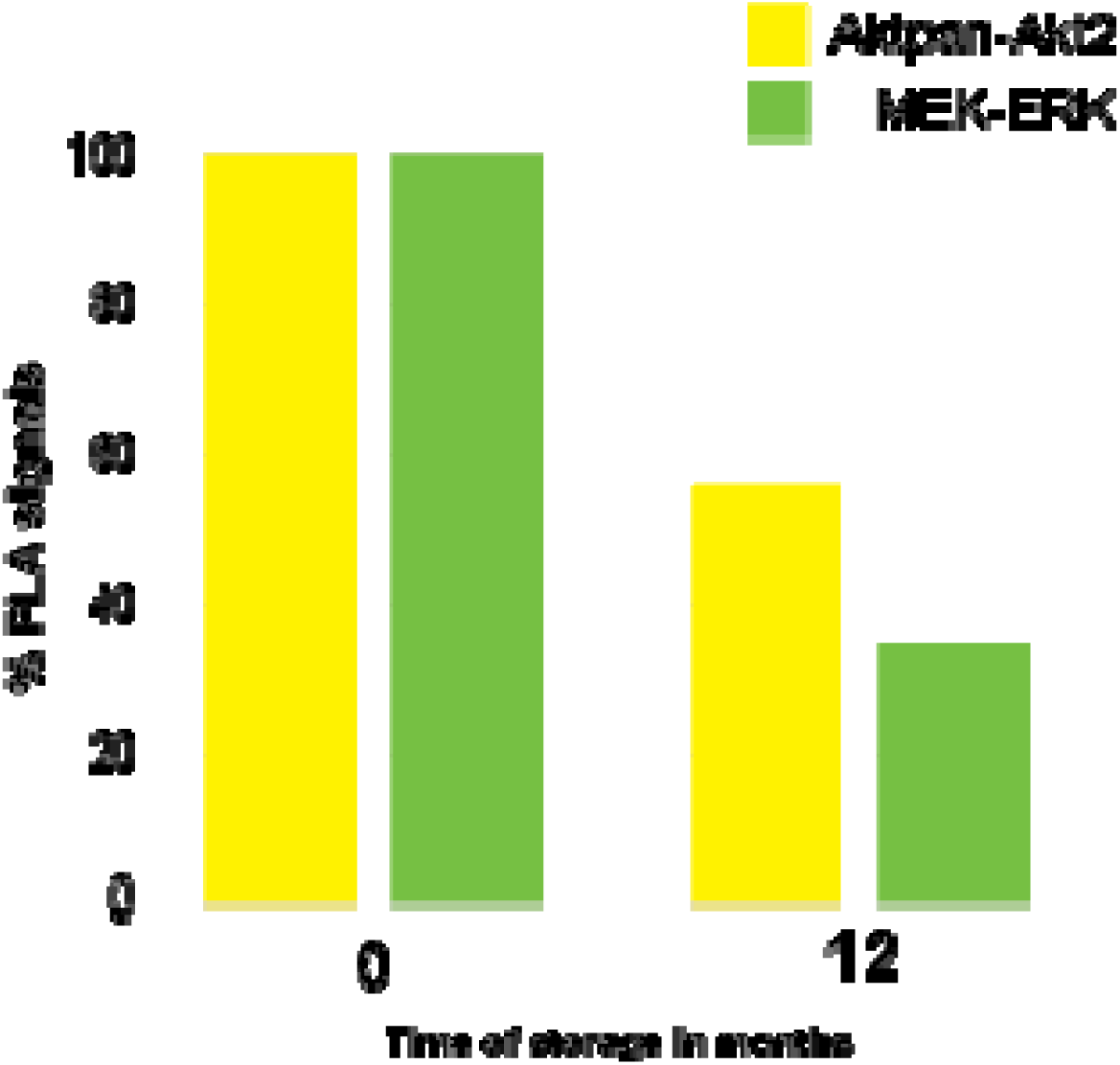
The figure shows SKBR-3 cells stored in -20C, from 0 to 12 months. The yellow bar represents the Aktpan-Akt2 protein interactions and the green bar represents the MEK-ERK protein interactions. The left pair of bars shows cells prepared and analyzed at time point zero and the pair to the right shows SKBR-3 cells analyzed 12 months later. SKBR-3 cells, which were freshly prepared was generated to be the optimal storage condition and represent 100% of the potential identified dots. The other bars show the relative decrease percentage of identified PLA-dots.

**Figure 5.**
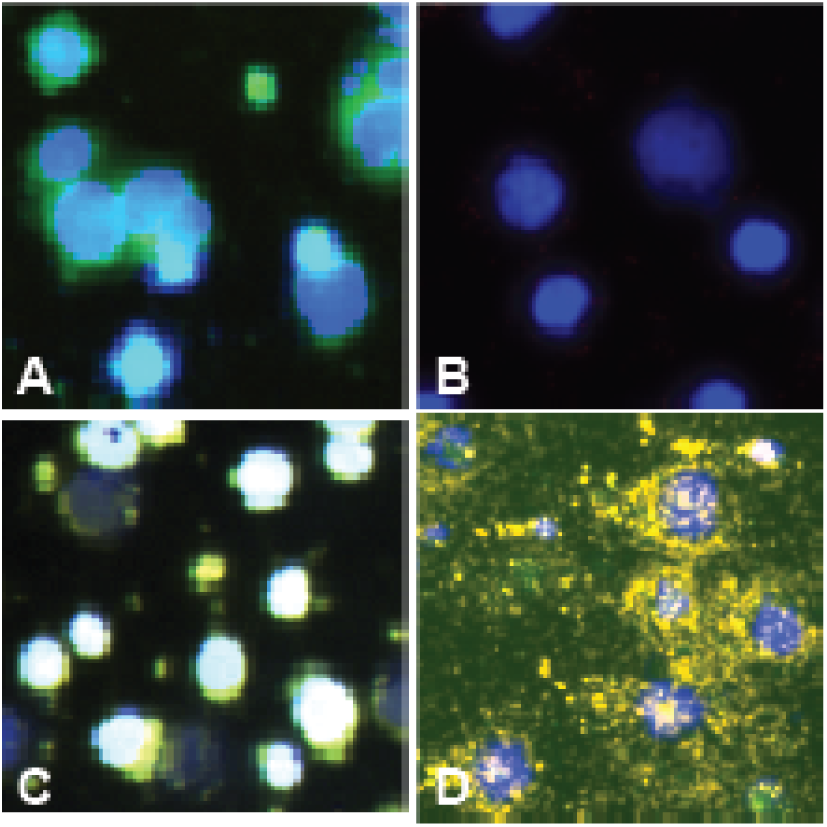
Representative fluorescence microscopy images demonstrating the effects of staining and storage temperature on isPLA signals and autofluorescence. A, MGG-stained GM12878 cells stored at RT, showing increased autofluorescence and few or no detectable isPLA signals. B, Negative control for GM12878 cells. C, Unstained Jurkat cells stored in RT, showing autofluorescence and no clearly detectable specific isPLA signals. D, Unstained GM12878 cells stored at-20 °C, The nuclei were stained with Hoechst. Aktpan-Akt2 and MEK-ERK interaction were detected by using Cy3- and FITC488-labelled probes, respectively. Original magnification 20x.

### 3.3 Morphological assessment of stored cytological specimens

Unstained GM12878 cells stored at RT exhibit virtually no detectable isPLA signals despite preserved cellular morphology, Figure. 5. In contrast, unstained cells stored at -20°C demonstrated abundant intracellular fluorescence signals corresponding to preserved protein interactions.

Slides stained with MGG and subsequently stored at RT exhibit increased autofluorescence together with an almost complete absence of detectable isPLA signals.

Similarly, SKBR-3 slides stored for one year at -20°C displayed fewer detectable intracellular protein interactions than newly prepared slides, Figure 6, consistent with a gradual reduction in protein detectability during prolonged storage.

**Figure 6.**
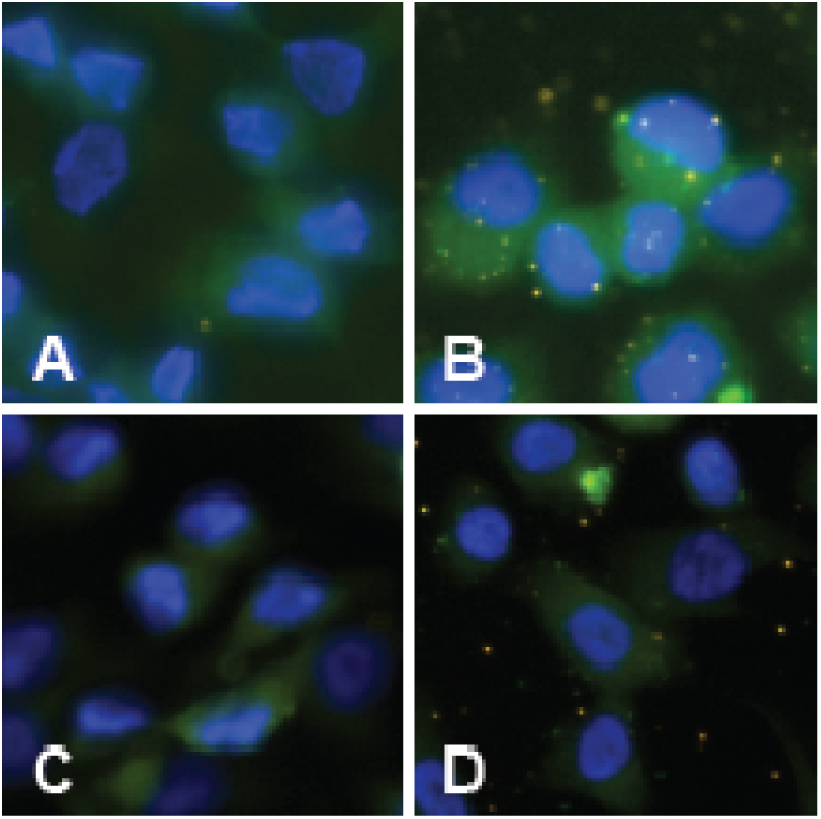
Representative fluorescence microscopy images showing reduced isPLA signal detection in SKBR-3. A, negative control at baseline. B, Positive control analyzed at baseline. C, negative control analyzed after 12 months of storage. D, Positive control analyzed efter 12 months of storage, showing reduced Aktpan-akt2 and MEK-ERK signals. Nuclei was stained with Hoechst. Interactions were detected using cy3- and FITC488 labelled probes, respectively.

### 3.4 Quantitative image analysis

Automated image analysis using CellProfiler successfully identified and segmented nuclei, cell boundaries and fluorescent isPLA signals based on DAPI, Cy3/554 and FITC channel. Quantification was based on approximately 10,000 cells per experimental set and demonstrated a consistent reduction in detectable protein interactions in specimen stored at RT compared with those stored at -20°C. The CellProfiler workflow enabled objective quantification of protein interactions and facilitated direct comparison between experimental conditions.

### 4.1 Discussion

The present study demonstrates that pre-analytical handling is a critical determinant of intracellular protein preservation in cytological specimens. Using isPLA, we show that storage temperature, staining procedures, and storage duration influence the detection of intracellular proteins and protein–protein interactions. Cytological slides stored at −20°C retained robust protein signals for prolonged periods, whereas storage at RT resulted in a marked reduction in detectable isPLA signals, accompanied by increased autofluorescence and loss of antigenicity. These findings emphasize that standardized pre-analytical handling is essential when archived cytological specimens are used for downstream protein analyses.

Biobanking has traditionally focused on preserving nucleic acids for genomic and transcriptomic analyses because DNA, and under appropriate conditions also RNA, can be reliably preserved in fixed cytological specimens. In contrast, proteins are considerably more susceptible to pre-analytical variables, including degradation, conformational alterations, epitope masking, and fixation-induced structural changes, all of which may affect protein detection and biological interpretation. Our findings are consistent with previous studies demonstrating that fixation and storage conditions substantially influence immunocytochemical staining and other molecular analyses of cytological specimens ^12,18,19^. However, comparatively little attention has been paid to the preservation of intracellular protein–protein interactions in archived cytological specimens, particularly in the context of proximity-based assays such as isPLA. Extending previous observations, our results demonstrate that standardized fixation and low-temperature storage preserve intracellular protein interactions and maintain robust isPLA signals for at least 24 months, whereas prolonged storage at RT results in a marked loss of signal intensity. We also noted a pronounced influence of storage temperature on the preservation of intracellular protein interactions. Slides stored at −20°C consistently maintained significantly higher numbers of detectable isPLA signals than those stored at room temperature. The profound signal loss observed during RT-storage may reflect progressive protein degradation, conformational alterations, oxidation, or reduced accessibility of protein epitopes. The increased autofluorescence observed in stained slides stored at room temperature may further impair signal detection by reducing the signal-to-noise ratio, thereby obscuring specific fluorescence signals and increasing the risk of false-negative results. Although the present study was not designed to determine the relative contribution of these individual mechanisms, the findings clearly demonstrate that storage temperature is a major determinant of the preservation of intracellular protein–protein interactions in archived cytological specimens.

May–Grünwald–Giemsa staining also influenced protein detectability. Nevertheless, our results indicate that stained cytological preparations remain suitable for protein analysis when staining, microscopic evaluation, and subsequent isPLA are performed within the same day. This observation is clinically relevant because routine cytology slides are frequently examined morphologically before additional analyses are considered, such as antibody-based immunocytochemistry or isPLA. WE note that isPLA provides cleaner and more specific results than IHC, which appears more diffuse and also more difficult to interpret, and isPLA can also reveal protein activity states as manifested in dynamic interactions and posttranslational modifications. Prolonged storage of stained slides resulted in substantial loss of detectable protein interactions, indicating that long-term storage of stained cytological preparations should be avoided when downstream protein analyses are anticipated.

The protein interactions targeted in the study demonstrated different preservation characteristics during prolonged storage in that the pan-Akt/Akt2 interaction remained detectable to a greater extent than the MEK–ERK interaction. Although the present study was not designed to investigate the underlying molecular mechanisms, these observations indicate that protein-specific stability should be considered when selecting biomarkers for retrospective analyses of archived cytological material. However, differential stability upon storage has also been reported in tissues ^20^. Further, one-year-old preparations of SKBR-3 cells exhibited fewer signals compared to those preserved for a shorter time. It is known that formaldehyde fixation could mask epitopes but maintain the morphology, and alcohol could degrade epitopes and lipids, but does not mask any epitopes ^21,22^. Further, different fixation depending on the biomarker is not the ideal protocol for multiplex assays, when both surface proteins, intracytoplasmic, intranuclear proteins, and possible phosphate proteins are included one panel^23^.

The combination of isPLA with automated image analysis strengthened the conclusions of this study, permitting extensive analyses. Approximately 39,000 cells were analyzed using CellProfiler, providing objective and reproducible quantification of intracellular protein interactions. Automated image analysis reduces observer-dependent variability and enables quantitative assessment of protein expression at the single-cell level. Such approaches are likely to become increasingly important as digital pathology and artificial intelligence are integrated into routine diagnostic workflows.

From a clinical perspective, minimally invasive cytology offers several advantages over tissue biopsy, including rapid sampling, low invasiveness, and the possibility of repeated longitudinal sampling. The ability to preserve intracellular protein integrity in archived cytological specimens expands the potential applications of cytology beyond conventional morphology and immunocytochemistry. Standardized cytology biobanking may therefore facilitate retrospective biomarker discovery, longitudinal monitoring of disease-associated signaling pathways, and future implementation of precision diagnostics based on single-cell protein profiling.

The present findings are also aligned with current international recommendations for biospecimen management, which emphasize that standardized pre-analytical handling is fundamental for preserving sample quality and ensuring reproducible molecular analyses. The International Society for Biological and Environmental Repositories (ISBER) Best Practices recommend standardized procedures for specimen collection, processing, storage, and documentation to maximize the long-term scientific value of bio banked material. The workflow proposed in the present study extends these principles to cytological specimens by providing a practical framework for preserving intracellular protein integrity and enabling reliable downstream single-cell protein analyses in archived cytology samples (21). The standardized cytology biobanking workflow developed and validated in the present study (Figure. 1) extends these principles to cytological specimens by preserving intracellular protein integrity and enabling reliable single-cell protein profiling in archived samples. This workflow provides a practical framework for retrospective biomarker discovery, translational pathology, and future implementation of precision diagnostics.

Several limitations should be acknowledged. First, this study was performed using standardized human cancer cell lines rather than patient-derived cytological specimens. Although standardized cell lines provide excellent experimental reproducibility and enable controlled evaluation of pre-analytical variables, future validation in archived patient-derived cytological specimens is warranted before clinical implementation. In addition, only a limited number of intracellular signaling pathways were evaluated, and additional biomarkers should be investigated in future studies. Furthermore, although long-term storage effects were examined for up to 24 months, dedicated biobank storage conditions may provide even greater protein stability than the routinely used laboratory freezer employed in this study. Future investigations should evaluate additional fixation methods, storage temperatures, and clinically relevant biomarkers across a broader range of cytological specimens.

In conclusion, standardized fixation, handling, and low-temperature storage preserve intracellular protein integrity and enable reliable detection of protein–protein interactions in archived cytological specimens. The standardized workflow presented here provides a practical framework for cytology biobanking and supports future applications in quantitative single-cell protein analysis, retrospective biomarker studies, and translational pathology.

This study demonstrates that intracellular protein interactions can be reliably visualized in cytology samples when appropriate handling and storage protocols are applied. Our results highlight the critical importance of using minimally invasive cytology sampling, combined with optimized fixation and storage protocols, to enable advanced molecular profiling and biomarker discovery at the single-cell level. Standardizing cytology sample handling will be key to expanding the use of cytological biobanking for personalized medicine and retrospective molecular studies.

## 5. Acknowledgement

Thanks to Professor Ulf Landegren for letting us use the in-house isPLA design.

## 7. Author Contributions

Conceptualization: Caroline Maria Dahlström (CMD), Jan-Michaél Hirsch (JMH),

Methodology: CMD, Liza Löf (LL).

Investigation: CMD.

Data curation: CMD.

Formal analysis: CMD, JMH.

Visualization: CMD.

Writing – original draft: CMD, JMH.

Writing – review & editing: CMD, JMH, LL. Supervision: JMH.

All authors read and approved of the final manuscript.

Funding acquisition: JMH

## References

1. Rodrigo JP, Rodríguez-Santamarta T, Corte D, et al. Hippo-YAP signaling activation and cross-talk with PI3K in oral cancer: A retrospective cohort study. Oral Dis. 2024;30(2):149–162. doi:10.1111/odi.14350

2. Abed FM, Dark MJ. Determining the utility of veterinary tissue archives for retrospective DNA analysis. PeerJ. 2016;4:e1996. doi:10.7717/peerj.1996

3. Celentano A, Glurich I, Borgnakke WS, Farah CS. World Workshop on Oral Medicine VII: Prognostic biomarkers in oral leukoplakia and proliferative verrucous leukoplakia-A systematic review of retrospective studies. Oral Dis. 2021;27(4):848–880. doi:10.1111/odi.13363

4. Alsarraf A, Kujan O, Farah CS. Liquid-based oral brush cytology in the diagnosis of oral leukoplakia using a modified Bethesda Cytology system. J Oral Pathol Med Off Publ Int Assoc Oral Pathol Am Acad Oral Pathol. 2018;47(9):887–894. doi:10.1111/jop.12759

5. Babshet M, Nandimath K, Pervatikar S, Naikmasur V. Efficacy of oral brush cytology in the evaluation of the oral premalignant and malignant lesions. J Cytol. 2011;28(4):165–172. doi:10.4103/0970-9371.86342

6. Chasma F, Pedr King R, Ker SY. Are there diagnostic alternatives to histopathology in detecting oral cancer? Evid Based Dent. 2022;23(1):24–25. doi:10.1038/s41432-022-0251-1

7. da Cunha Santos G, Schroder M, Zhu JB, et al. Minimizing delays in DNA retrieval: The “freezer method” for glass coverslip removal. Letter to the editor regarding comparative study of epidermal growth factor receptor mutation analysis on cytology smears and surgical pathology specimens from primary and metastatic lung carcinomas. Cancer Cytopathol. 2013;121(9):533. doi:10.1002/cncy.21306

8. Dejmek A, Zendehrokh N, Tomaszewska M, Edsjö A. Preparation of DNA from cytological material: effects of fixation, staining, and mounting medium on DNA yield and quality. Cancer Cytopathol. 2013;121(7):344–353. doi:10.1002/cncy.21276

9. Böning S, Schneider F, Huber AK, et al. Region of interest localization, tissue storage time, and antibody binding density-a technical note on the GeoMx® Digital Spatial Profiler. Immuno-Oncol Technol. 2024;23:100727. doi:10.1016/j.iotech.2024.100727

10. Colburn ME, Delaney MA, Anchor GC, Terio KA. Effect of formalin-fixation and paraffin-embedded tissue storage times on RNAscope in situ hybridization signal amplification. J Vet Diagn Investig Off Publ Am Assoc Vet Lab Diagn Inc. 2024;36(4):498–505. doi:10.1177/10406387241245777

11. Nam SK, Im J, Kwak Y, et al. Effects of fixation and storage of human tissue samples on nucleic Acid preservation. Korean J Pathol. 2014;48(1):36–42. doi:10.4132/KoreanJPathol.2014.48.1.36

12. Skoog L, Tani E. Immunocytochemistry: an indispensable technique in routine cytology. Cytopathol Off J Br Soc Clin Cytol. 2011;22(4):215–229. doi:10.1111/j.1365-2303.2011.00887.x

13. Kanber Y, Pusztaszeri M, Auger M. Immunocytochemistry for diagnostic cytopathology-A practical guide. Cytopathol Off J Br Soc Clin Cytol. 2021;32(5):562–587. doi:10.1111/cyt.12993

14. Shi Y, Yee-Chang M, Shi SR. Application of Immunohistochemistry in Cytology. Appl Immunohistochem Mol Morphol AIMM. 2023;31(7):459–466. doi:10.1097/PAI.0000000000001086

15. Söderberg O, Leuchowius KJ, Gullberg M, et al. Characterizing proteins and their interactions in cells and tissues using the in situ proximity ligation assay. Methods San Diego Calif. 2008;45(3):227–232. doi:10.1016/j.ymeth.2008.06.014

16. Alam MS. Proximity Ligation Assay (PLA). Curr Protoc Immunol. 2018;123(1):e58. doi:10.1002/cpim.58

17. Leuchowius KJ, Weibrecht I, Söderberg O. In situ proximity ligation assay for microscopy and flow cytometry. Curr Protoc Cytom. 2011;Chapter 9:Unit 9.36. doi:10.1002/0471142956.cy0936s56

18. Pinheiro C, Roque R, Adriano A, et al. Optimization of immunocytochemistry in cytology: comparison of two protocols for fixation and preservation on cytospin and smear preparations. Cytopathol Off J Br Soc Clin Cytol. 2015;26(1):38–43. doi:10.1111/cyt.12156

19. Mirlacher M, Kasper M, Storz M, et al. Influence of slide aging on results of translational research studies using immunohistochemistry. Mod Pathol Off J U S Can Acad Pathol Inc. 2004;17(11):1414–1420. doi:10.1038/modpathol.3800208

20. Crowe AR, Yue W. Semi-quantitative Determination of Protein Expression using Immunohistochemistry Staining and Analysis: An Integrated Protocol. Bio-Protoc. 2019;9(24):e3465. doi:10.21769/BioProtoc.3465

21. Cizkova K, Flodrova P, Baranova R, Malohlava J, Lacey M, Tauber Z. Beneficial Effect of Heat-induced Antigen Retrieval in Immunocytochemical Detection of Intracellular Antigens in Alcohol-fixed Cell Samples. Appl Immunohistochem Mol Morphol AIMM. 2020;28(2):166–174. doi:10.1097/PAI.0000000000000689

22. Unhale SA, Skubitz APN, Solomon R, Hubel A. Stabilization of tissue specimens for pathological examination and biomedical research. Biopreservation Biobanking. 2012;10(6):493–500. doi:10.1089/bio.2012.0031

23. Sakabe N, Maruyama S, Ito C, et al. Effect of liquid-based cytology fixing solution on immunocytochemistry: Efficacy of antigen retrieval in cytologic specimens. Diagn Cytopathol. 2023;51(9):546–553. doi:10.1002/dc.25178

